# Single cell lineage dynamics of the endosymbiotic cell type in a soft coral *Xenia* species

**DOI:** 10.1101/2019.12.12.874602

**Authors:** Minjie Hu, Xiaobin Zheng, Chen-Ming Fan, Yixian Zheng

## Abstract

Many hard and soft corals harbor algae for photosynthesis. The algae live inside coral cells in a specialized membrane compartment called symbiosome, which shares the photosynthetically fixed carbon with coral host cells, while host cells provide inorganic carbon for photosynthesis^1^. This endosymbiotic relationship is critical for corals, but increased environmental stresses are causing corals to expel their endosymbiotic algae, i.e. coral bleaching, leading to coral death and degradation of marine ecosystem^2^. To date, the molecular pathways that orchestrate algal recognition, uptake, and maintenance in coral cells remain poorly understood. We report chromosome-level genome assembly of a fast-growing soft coral, *Xenia* species (*sp.*)^3^, and its use as a model to decipher the coral-algae endosymbiosis. Single cell RNA-sequencing (scRNA-seq) identified 13 cell types, including gastrodermis and cnidocytes, in *Xenia sp*. Importantly, we identified the endosymbiotic cell type that expresses a unique set of genes implicated in the recognition, phagocytosis/endocytosis, maintenance of algae, and host coral cell immune modulation. By applying scRNA-seq to investigate algal uptake in our new *Xenia sp*.. regeneration model, we uncovered a dynamic lineage progression from endosymbiotic progenitor state to mature endosymbiotic and post-endosymbiotic cell states. The evolutionarily conserved genes associated with the endosymbiotic process reported herein open the door to decipher common principles by which different corals uptake and expel their endosymbionts. Our study demonstrates the potential of single cell analyses to examine the similarities and differences of the endosymbiotic lifestyle among different coral species.

---

Similar to other endosymbiotic cnidarian, corals take up the *Symbiodiniaceae* family of dinoflagellate algae into their gastrodermis through feeding. Some cells in the gastrodermis, lining the digestive tract, appear to have the ability to recognize certain types of algae. Through phagocytosis and by modulating host immune responses, the correct alga type is preserved and enclosed by endomembranes to form symbiosomes in coral cells^1^. The symbiosome membrane is believed to contain transporters that mediate nutrient exchange between the algae and host cells^4^. In recent years, many comparative transcriptome analyses were performed on whole organisms to identify gene expression changes before and after algae colonization or bleaching using different cnidarian species. This led to the identification of genes whose up- or down-regulation could contribute to endosymbiosis. Comparative genomic and transcriptomic information in endosymbiotic and non-endosymbiotic cnidaria species were also used to search for genes that may have evolved to mediate recognition or endocytosis of *Symbiodiniaceae*^5–8^. These approaches, however, do not differentiate whether the altered genes are expressed in the host endosymbiotic cells or other cell types without additional criteria. Protein inhibition or activation studies were also used to suggest the host proteins containing C-type lectin domains, scavenger receptor domains, or thrombospondin type 1 repeats to be involved in algal uptake and immunosuppression^9–11^. The broad expression and function of these proteins coupled with potential off-target effects of inhibitors greatly limit data interpretation. Therefore, a systematic description of genes and pathways selectively expressed in the host endosymbiotic cells is much needed to begin to understand the potential regulatory mechanisms underlying the entry, establishment, and possibly expulsion of *Symbiodiniaceae*.

To enable systematic study of coral biology, we reason that it is important to first define gene expression characteristics of each cell type in corals because it would offer an opportunity to identify the molecular signature of the cell type that performs endosymbiosis. We focused on a soft coral pulsing *Xenia sp*. (Fig. 1a, b, Supplementary Video 1) that grows in a laboratory aquarium rapidly. Using Illumina short-read and Nanopore long-read sequencing (Extended data Tables 1, 2), we assembled the *Xenia* genome into 556 high quality contigs. Applying chromosome conformation capture (Hi-C)^12^, we further assembled these contigs into 168 scaffolds with the longest 15 contain 92.5% of the assembled genome of 222,699,500 bp, which is similar to GenomeScope estimated genome size (Fig. 1c, Extended Data Fig. 1). The *Xenia* genome has by far the longest scaffold length and thus most contiguous assembly compared to the published cnidarian genomes (Fig. 1d). Genome annotation using several bulk RNA-sequencing (RNA-seq) data showed that *Xenia sp*. has similar number of genes as other cnidaria (Extended Data Tables 3, 4). Consistent with previous phylogenetic analysis^13^, soft corals *Xenia sp*. and *Dendronephthya gigantean* grouped together, they branched later than the freshwater *Hydra* from the stony corals, and they are more distantly related to the stony corals than *Nematostella* and *Aiptasia* (Fig. 1e).

**Fig. 1.**
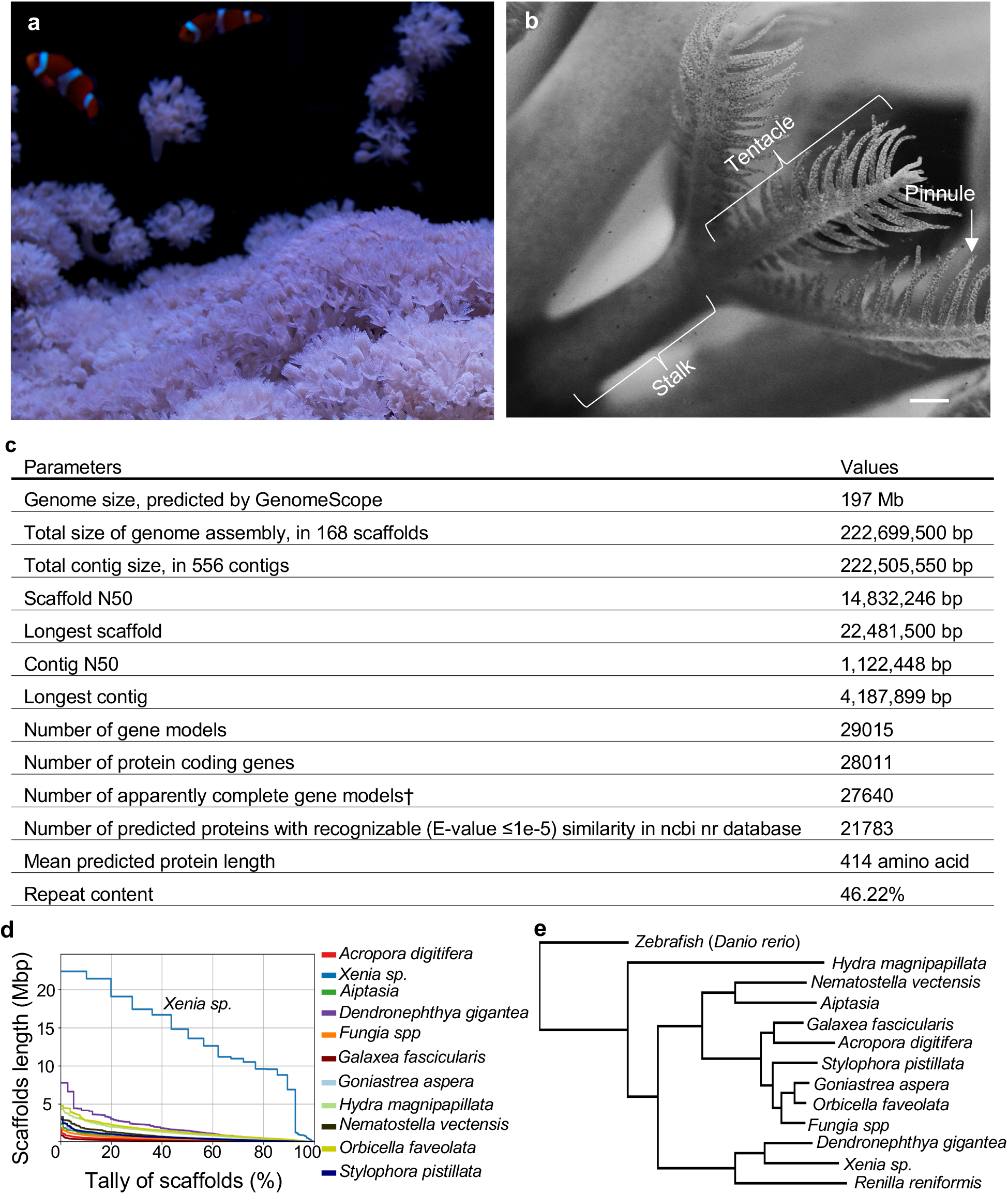
High quality genome assembly for *Xenia sp*. **a**, *Xenia sp*. grown in a laboratory aquarium. **b**, An enlarged view of a *Xenia sp*. polyp with its main sub-structures indicated. Scale bar, 1 mm. **c**, A summary of *Xenia sp*. genome assembly and gene annotation. †, genes encoding protein sequences with apparent in frame start and stop codons. **d**, Comparisons of the assembled scaffold lengths (y-axis) and tallies (x-axis) of 11 sequenced cnidarians including *Xenia sp*. **e**, Evolutionary comparisons of *Xenia sp*. with other cnidarians as indicated. Zebrafish was used as an outgroup. The phylogenetic branch points were assigned with 100% confidence.

We next performed single cell RNA-seq (scRNA-seq)^14^ of whole polyps, stalks, or tentacles using the 10xGenomics platform. By t-distributed stochastic neighbor embedding (t-SNE)^15^, we grouped the high-quality single cell transcriptomes covering 19,976 genes into 13 cell clusters with distinct gene expression patterns (Fig. 2a, b, Extended Data Fig. 2a, Supplementary Table 1). For validation, we attempted to identify two previously characterized cnidarian cell types, cnidocytes for prey capture/defense and gasterodermis cells. We found that cluster 12 cells express cnidocyte marker genes, *Minicollagens* and *Nematogalectins*^16–18^ (Fig. 2c). Further analysis of cluster 12 single-cell transcriptomes revealed two sub-clusters (Fig. 2d, Extend Data Fig. 2b). *Minicollagens* are expressed in both sub-clusters, whereas *Nematogalectins* are perferentially expressed in one sub-cluster (Fig. 2e and Extended Data Fig. 2c). RNA *in situ* hybridization (ISH) confirmed the expression of a *Nemetogalectin* to be more restricted than the *Minicollagen* in *Xenia* pinnules (Fig. 2f, g). Interestingly, *Xenia* cells in clusters 2, 8, and 13 express genes encoding collagens and proteases (Fig. 2h) shown to be enriched in gastrodermis of *Nematostella*^16^. RNA ISH for *Collagen 6* (expressed by clusters 2 and 8), *Astacin-like metalloendopeptidase2* and another gene Xe_003623 (both expressed by all 3 clusters) showed their presence in the gastrodermis (Fig. 2h-j, Extended Data Fig. 2d-h). Thus, our clustering analyses has identified different types of cnidocytes and gastrodermis cells in *Xenia*.

**Fig. 2.**
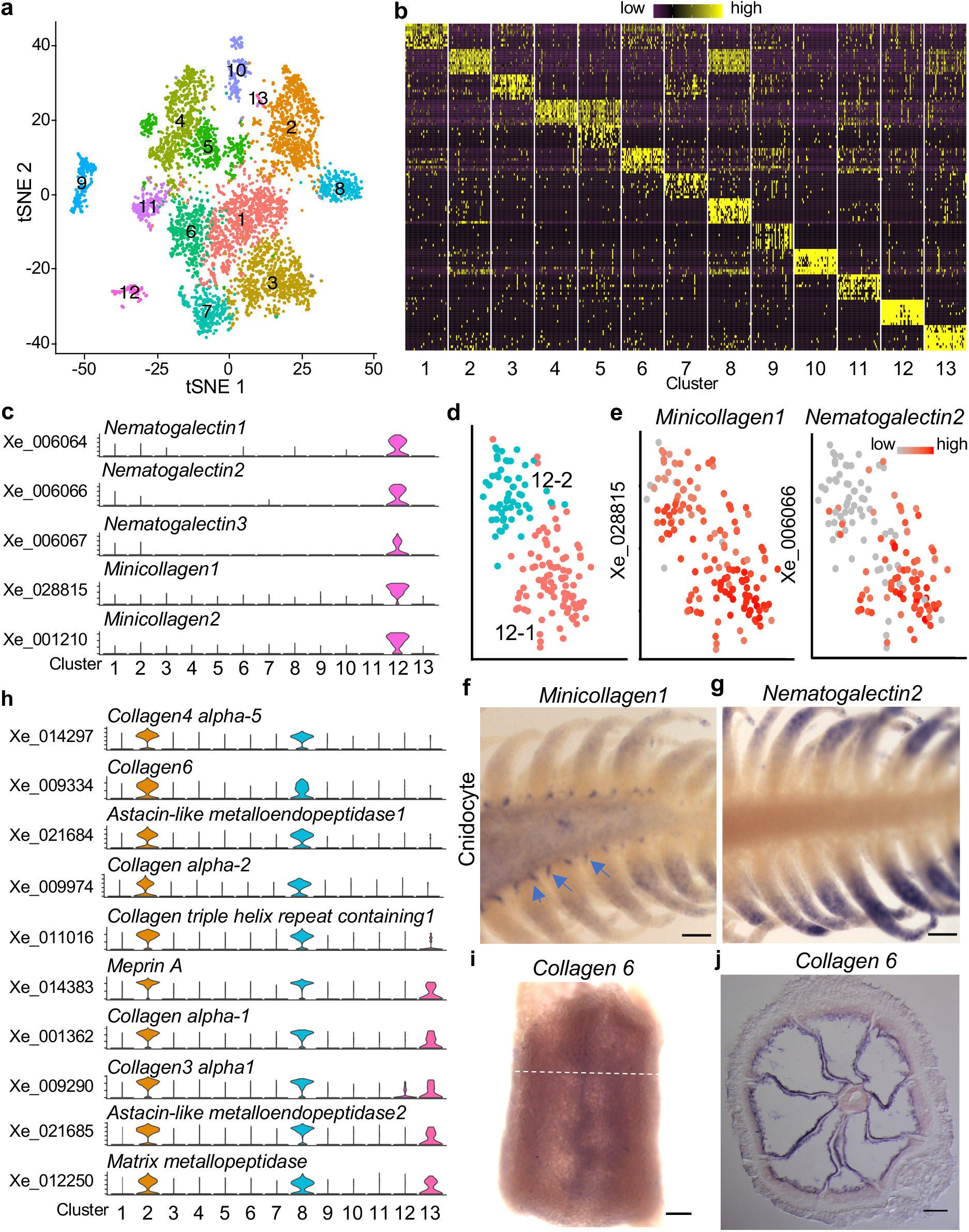
scRNA-seq transcriptomes delineate 13 cell types in *Xenia sp*. **a**, Transcriptomes of individual *Xenia sp*. cells obtained by scRNA-seq were grouped into 13 clusters (color coded) and presented in the tSNE space. Each colored dot represents one cell. **b**, Gene expression heatmap (scale on the top) for the top 10 gene markers defining each cluster. Each column and row represent one cell cluster and gene, respectively. 40 cells were randomly selected from each of the 13 cell clusters for plotting. **c**, Expression profiles of the indicated marker genes defining cluster 12. **d**, Cluster 12 cells is sub-divided into two populations (12-1 and 12-2, color-coded) and displayed in a tSNE space. Each colored dot represents a cell. **e**, Expression levels (scale to top right) of two cluster 12 markers, *Minicollagen1* and *Nematogalectin2*, are shown in the tSNE plot. **f, g**, Whole mount RNA *in situ* hybridization of *Minicollagen1* (**f**) and *Nematogalectin2* (**g**) showing their expression in tentacles. Arrows indicate the expression of *Minicollagen1* at the base of pinnules. **h**, Expression profiles of marker genes enriched in clusters 2, 8, and 13. **i, j**, RNA *in situ* hybridization of *Collagen 6*. Whole mount view of the stalk in **i** and cross section image in **j**. The white dashed line in **i** indicates the cross section level in **j**. Scale bars, 100 μm (**f, g, j**), 150 μm (i).

To identify the cell type for endosymbiosis, we took advantage of the auto-fluorescence of *Symbiodinaceae*. Using fluorescence-activated cell sorting (FACS), we separate the algae-containing and algae-free *Xenia* cells (Fig. 3a, b, Extended Data Fig. 3a-d) and performed bulk RNA-seq of these cell populations (Supplementary Table 2). By comparing these bulk cell transcriptomes with genes expressed in each cluster, we found that cluster 13 cells exhibited the highest similarity with the algae-containing cells (Fig. 3c, d, Supplementary Table 3, 4). RNAscope ISH for two cluster 13 marker genes, one encoding a protein with <u>Le</u>ctin and Kazal <u>Pr</u>otease <u>in</u>hibitor domains (LePin) and the other encoding Granulin 1, showed expression of both genes in algae-containing gastrodermis cells (Fig. 3e, Extended Data Fig. 3e, f). Thus, of the 3 gastrodermis cell types, cluster 13 cells play a unique role in endosymbiosis.

**Fig. 3.**
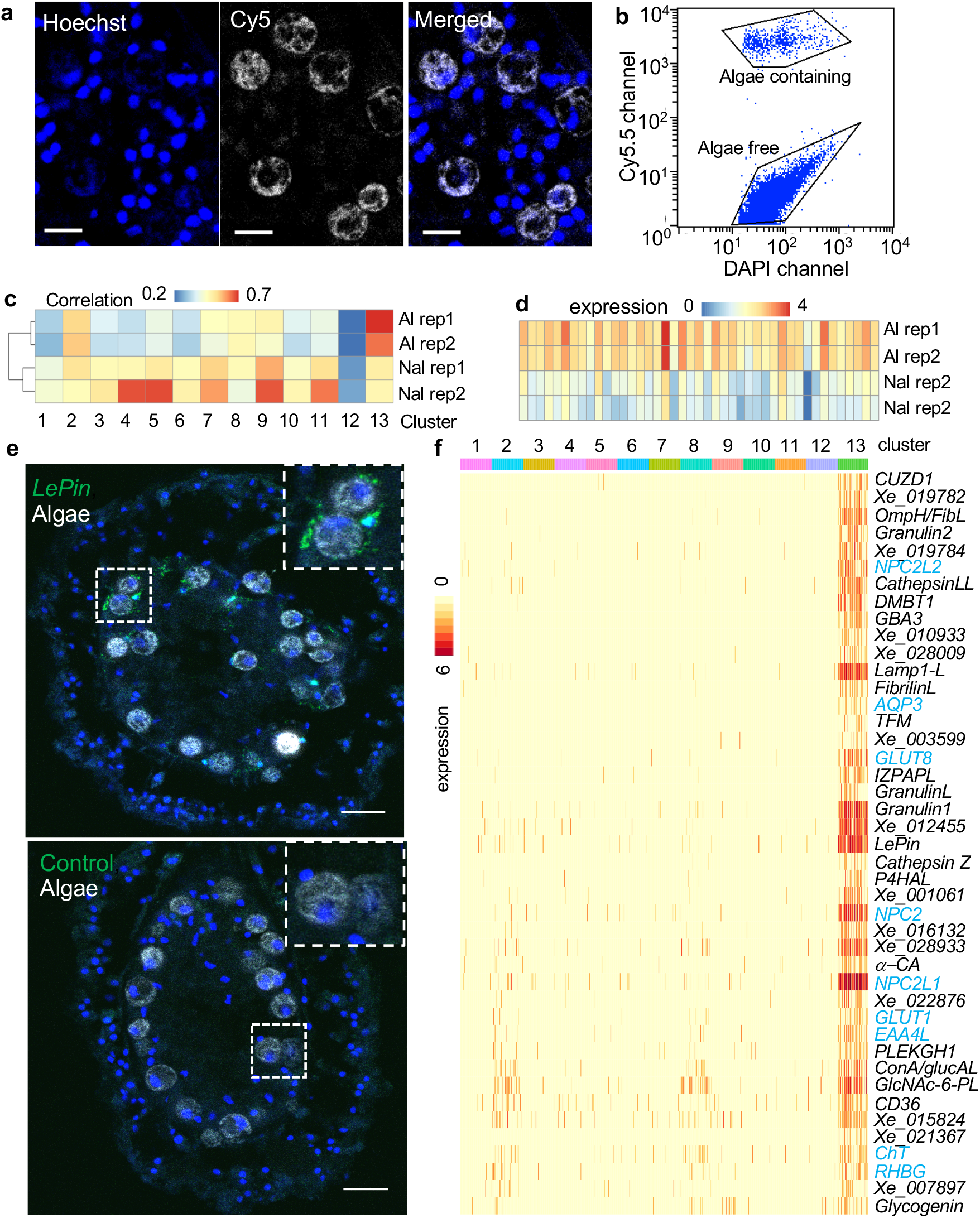
Identification of genes specifically expressed in *Xenia sp*. endosymbiotic cells. **a**, The endosymbiotic algae in *Xenia* display autofluorescence in the far-red channel. A cross section of *Xenia* with *Xenia* and algal nuclei stained by Hoechst in blue and algae autofluorescence in white. **b**, A FACS profile of dissociated live *Xenia* cells using Cy5.5 and DAPI channels. **c**, Pearson correlation of gene expression between the scRNA-seq data of 13 cell clusters and the bulk RNA-seq data of two biological replicates of FACS-isolated algae-containing (Al rep1 and Al rep2) and algae-free (Nal rep1 and Nal rep2) cells. **d**, Heatmap showing the expression levels of the top 43 markers genes for cluster 13 cells in FACS-sorted algae-containing and algae-free cells. **e**, Ultra-sensitive fluorescence RNA *in situ* hybridization by RNAscope probing for *LePin* (green, top) and control (bottom). A *Xenia* cell with algae and *LePin* expression is enlarged to the top right. Algae autofluorescence is in white. Hoechst staining of all nuclei are in blue. Scale bars, 10 μm. **f**, Heatmap showing the enrichment levels of the cluster 13’s top 43 marker genes in all 13 cell clusters. Transporters are highlighted in blue.

Among the top 43 marker genes highly enriched in cluster 13 endosymbiotic cells, 30 encode proteins with domains of known or predicted functions, including receptors, extracellular matrix proteins, immune response regulators, phagocytosis/endocytosis, or nutrient transports (Fig. 3f, Extended Data Fig. 4, Supplementary Table 3). Three proteins, encoded by *CD36, DMTB1*, and *CUZD1*, contain CD36 or scavenger receptor (SR) domains known to recognize a wide range of microbial surface ligands and mediate their phagocytosis, while modulating host innate immune response (Extended Data Fig. 4a)^10,19,20^. Among them, CUZD1 is least understood, but it is similar to DMBT1 in domain organization. Like CD36, DMBT1 functions in pattern-recognition of microbes. DMBT1 is expressed on the surface of mammalian gastrointestinal tracts where it recognizes poly-sulfated and poly-phosphorylated ligands on microbes, represses inflammatory response, and regulates gastro-intestinal cell differentiation^21^. *LePin* and *Granulin1*, used in ISH, have homologs in *Aiptasia*, stony and soft corals. Since LePin has an N-terminal signal peptide followed by multiple domains including H- and C-type lectins and Kazal type serine protease inhibitor (Extended Data Fig. 4b), it may confer selectivity for *Symbiodiniaceae*. *Granulin1* is one of three *Granulin*-related genes expressed selectively in endosymbiotic cells and in mammals Granulins were shown to modulate immune response^22^.

Phagocytosis of *Symbiodiniaceae* by the similar sized gastrodermis cells requires substantial host cell expansion, but genes regulating the expansion have remained unknown. Among the endosymbitoic marker genes, we found *Plekhg5*, which encodes a highly conserved RhoGEF (Extended Data Fig. 4c). The *Xenopus* Plekhg5 localizes to the apical membrane of epithelial cells and recruits actomyosin to induce cell elongation and apical constriction^23^. Thus, *Plekhg5* is a prime candidate in regulating the apical membrane extension of endosymbiotic cells to engulf *Symbiodiniaceae* during early stages of phagocytosis. Upon phagocytosis, *Symbiodiniaceae* are enclosed by the host membrane to form symbiosomes^24^. Although the symbiosome compartment is acidified similarly to lysosomes^25^, it is unclear whether specific genes are involved in symbiosome formed. *Xenia sp*. has two genes encoding lysosome-associated membrane glycoproteins (Lamp) that are more similar to the characterized Lamp1 than Lamp2. The *Xenia Lamp1-L* encodes a larger protein and is an endosymbiotic marker gene, whereas the *Lamp1-S* encodes a smaller Lamp1 and is uniformly expressed across cell clusters (Extended Data Fig. 4d, e). Since Lamps are known to regulate phagocytosis, endocytosis, lipid transport, and autophagy^26^, Lamp1-L may regulate symbiosome formation and/or function. Several endosymbiotic marker genes encode enzymes that may promote the establishment of endosymbiosis or facilitating nutrient exchanges between algae and the host. For example, *α-CA* is a predicted transmembrane carbonic anhydrase that may concentrate CO_2_ for photosynthesis by the algae. There are also nine genes potentially participate in nutrient exchanges as they encode highly conserved transporters for sugar, amino acid, ammonium, water, cholesterol, or choline (Fig. 3f).

To better understand the lineage sequence and temporal dynamics of the *Xenia* endosymbiotic cells, we developed a *Xenia* regeneration model. We surgically cut away all tentacles from *Xenia* polyps and found the stalks regenerated all tentacles in several days (Fig. 4a). Individual tentacles also regenerated into full polyps but requiring a longer time (not shown). BrdU labeling showed that the proliferated BrdU^+^ cells in the gastrodermis began to take up algae at day 4 of regeneration (Extended Data Fig. 5a, b). We performed scRNA-seq of the regenerating stalks and pooled the data with the scRNA-seq samples described above, followed by clustering them into 13 cell types. Nearly all cell clusters from the non-regeneration samples matched the clusters in the pooled samples and 100% of cells in the endosymbiotic cell cluster 13 in non-regeneration samples are found in the endosymbiotic cluster 13 in the pooled samples (Extended Data Fig. 5c). This allowed us to define endosymbiotic cells in the regenerative sample.

**Fig. 4.**
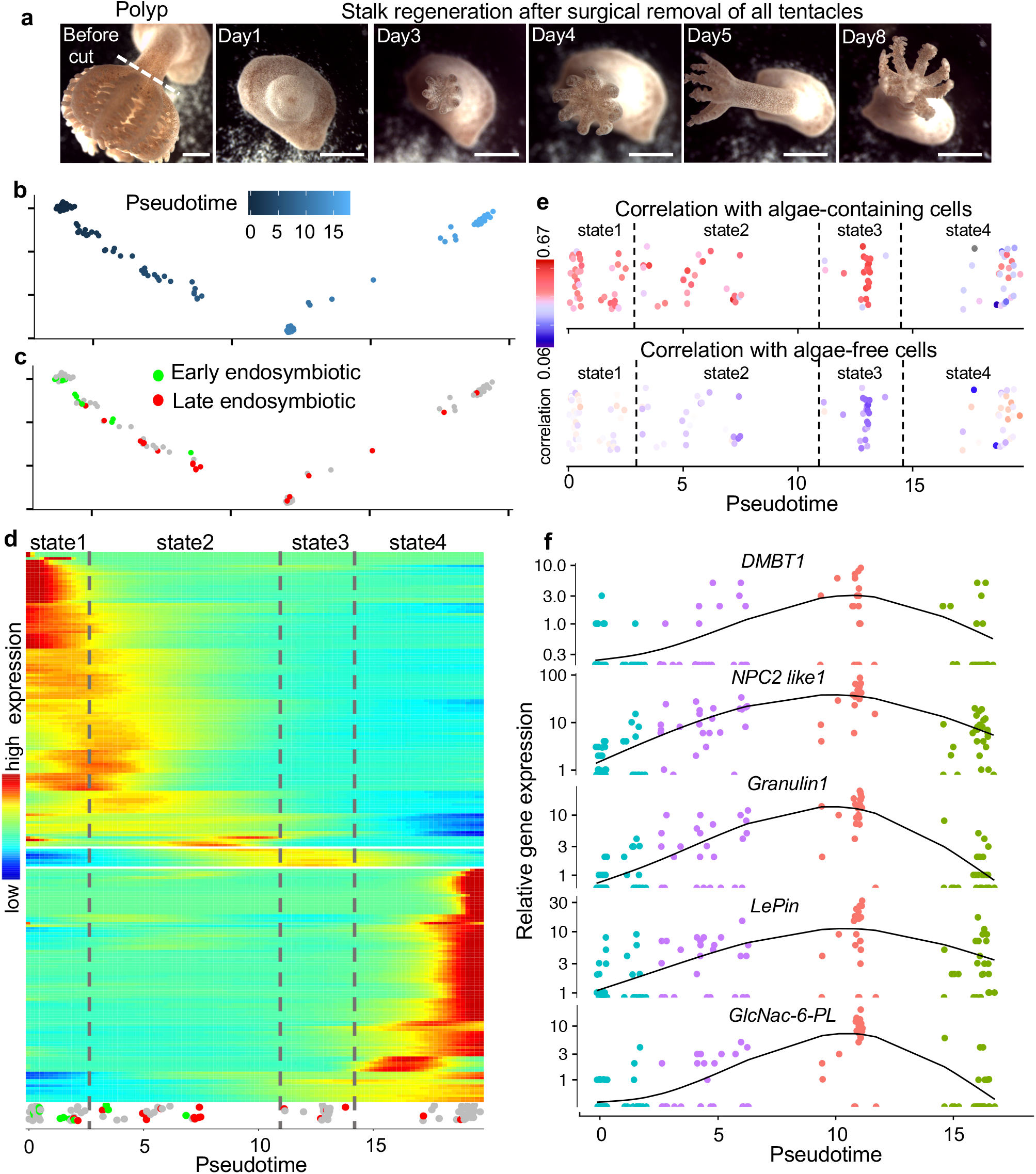
Dynamic lineage progression of endosymbiotic cells. **a**, An example *Xenia sp*. polyp shown in the first image is surgically cut at the white dashed line to remove all the tentacles. A surgically cut stalk is shown to regenerate in successive days as indicated. Scale bars, 1 mm. **b,** Pseudotime trajectory of all endosymbiotic cells (a dot represents a cell) identified in regeneration and non-regeneration scRNA-seq datasets. The pseudotime indicator is shown at the top. **c**, Distribution of early (green) and late (red) lineage cells predicted by velocyto based on scRNA-seq of the regenerating sample is highlighted on the same pseudotime plot shown in (**b**). **d**, Heatmap for gene expression levels along pseudotime. The endosymbiotic cells (as in **c**) used to model gene expression along the pseudotime line are shown at the bottom for reference; color key for gene expression level is shown to the left. Four cell states (state1-4, separated by dashed lines and indicated at the top) are defined by differential gene expression. **e**, Gene expression correlation between individual cells (shown as dots) in 4 cell states and the transcriptomes of the FACS-isolated algae-containing (top) or algae-free (bottom) cells; The heatmap key at the left shows Pearson correlation coefficient. **f**, Expression dynamics of endosymbiotic marker genes in the pseudotime through 4 cell states: cyan, purple, flesh, and green, represent state1, 2, 3, 4 cells (each dot represents a cell), respectively.

We performed pseudotemporal ordering of the endosymbiotic *Xenia* cells by Monocle 2^14^ (Fig. 4b), which uses reversed graph embedding to construct a principal curve that passes through the middle of the cells on the tSNE space. Because this trajectory analysis does not provide a direction of cell state progression, we used velocyto^27^ to determine the directionality of lineage progression of endosymbiotic cells in the regenerating sample. Velocyto calculates RNA velocity by comparing the number of un-spliced and spliced reads, which measure the expected change of gene expression in the near future, thereby providing the directionality of cell state change. This allowed the identification of early and late stages of endosymbiotic cells in the regenerating sample (Extended Data Fig. 5d). The cell trajectory showed that the early (green) and late (red) stage cells are mapped to the early and late pseudotime, respectively (Fig. 4c), revealing the pseudotime represents actual lineage progression. Gene expression modeling revealed significant changes along the pseudotime. Further hierarchical clustering revealed distinct gene expression patterns, which helped us to define four endosymbiotic cell states (Fig. 4d, Supplementary Table 5).

To further understand the cells in the four states, we compared single cell transcriptomes against transcriptomes from the bulk RNA-seq of FACS-sorted algae-containing or algae-free cells and plotted the expression correlation along the pseudotime. State3 cells showed the strongest correlation with the algae-containing cells followed by state1 and 2, whereas state4 cells showed the least correlation (Fig. 4e). Thus, state3 represents the mature agal-containing cells. Moreover, whereas state3 cells showed least correlation with algae-free cells, state1 and 4 cells showed similar correlations with algae-containing and algae-free cells (Fig. 4e). These suggest that state1 cells are pre-endosymbiotic progenitors that can transition through state2 to become state3 mature algae-containing cells, while state4 represents post-endosymbiotic cells. Consistent with this, the endosymbiotic marker genes we identified showed a gradual upregulation along the psuedotime from state1 through state2 to peak at state3, followed by a down regulation in state4 (Fig. 4f).

Analyzing the differentially expressed genes in each state suggests their roles in regulating endosymbiotic cell lineage development. For example, the state1 cells express two genes encoding a Secreted Frizzled-Related Protein (SFRP) and Elk1. SFRP modulates Wnt signaling during development and differentiation, whereas Elk1 is known to regulate cell proliferation and differentiation^28,29^. State1 cells also express a G-protein coupled receptor Mth with an N-terminal extracellular domain, which should allow the creation of specific antibodies that recognize these cells for live-sorting by FACS to aid future characterization of cells at this state. State1 and 2 cells express genes encoding thrombospondin type 1 repeats (TSR). Since TSR binds to CD36, which can convert TGFβ to active form to regulate immune tolerance^30^, the TSR-containing genes expressed in state1/2 cells may directly suppress immune response of host endosymbiotic cells during *Symbiodiniaceae* recognition and uptake. Our analyses suggest state4 cells as post-endosymbiotic. Further studies of the genes defining state4 cells should clarify if these cells lose algae due to aging or stress, and if they share features of bleached endosymbiotic cells.

Here we demonstrate the power of advanced genomic and bioinformatic tools in studying coral biology. Although we focused on studying the endosymbiotic cell lineage, the regenerative processes for the other cell types can be similarly comprehended as more information becomes available. We reveal 4 distinct *Xenia* endosymbiotic cell states and propose a lineage progression from state1 progenitors through state2 cells engaging in algae recognition to state3 mature endosymbiotic cells followed by state4 post-endosymbiotic status. Since state4 transcriptome has lowest expression of endosymbiotic marker genes among all four states, we speculate that these post-endosymbiotic cells represent a terminal state, which is either no-longer competent for algae re-uptake or can only regain endosymbiotic competence after a long recovery. The presence of state4 could offer new insights into why coral bleaching is devastating because bleached endosymbiotic cells may be similar to the state4 cells that cannot re-uptake or require a long time to recover and re-uptake algae. It will be interesting to test if efficient recovery from bleaching relies on state1 progenitors to make new endosymbiosis-competent cells. It is also feasible to further test whether forced regeneration by fragmenting bleached corals can stimulate state1 progenitor expansion and endosymbiosis restoration.

## Acknowledgement

We thank Fred Tan and Allison Pinder for assistance with all the sequencing. Yun Bai for assistance with cell sorting. Lynne Hugendubler and Mike Watts for maintaining the coral tank. Supported by Carnegie Institution for Science (YZ, CMF), NIH/NIGMS GM106023 (YZ), GM110151 (YZ), NIH/NIAR AR060042 (CMF), and AR071976 (CMF).

## Author contributions

CMF and YZ conceived and supervised the project. MH, CMF, and YZ designed experiments. MH performed the experiments. MH and XZ analyzed the data. MH, XZ, CMF, and XZ interpreted the data and wrote the manuscript.

---

**Extended Data Fig. 1.**
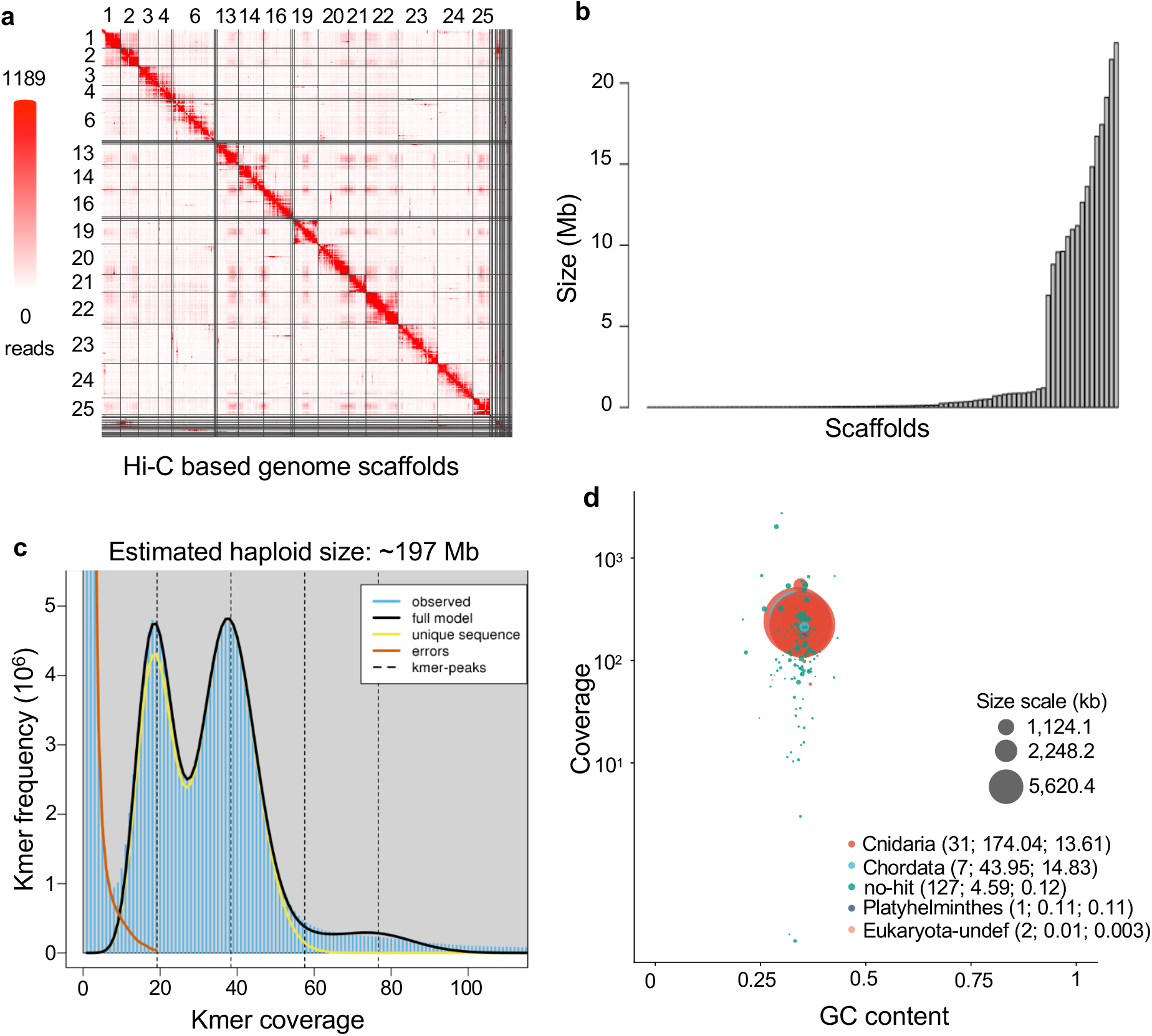
Additional genome assembly data. **a**, Hi-C based *Xenia sp*. genome assembly. The scaffolds are separated by grids demarcated by black lines. The numbers for the 15 longest scaffolds out of the total 168 are shown. **b**, Size distribution of *Xenia sp*. genome scaffolds with each bar on the x-axis representing a scaffold. **c**, *Xenia sp*. genome is predicted to be diploid, as expected, with a haploid genome size of ~197 Mb based on GenomeScope analysis of Illumina short reads. **d**, Contamination analysis by BlobTools revealed a similar GC content and genomic coverage across most scaffolds. Each colored circle in the graph represents a scaffold. Larger circles have longer scaffold lengths (see the three grey circles for length scale used in the plot). The color codes represent the closest species group that have the highest sequence similarities to the *Xenia sp*. scaffolds (the first number in each parenthesis shows the *Xenia* scaffold number followed by the combined length of the scaffolds and scaffold N50 in Mb).

**Extended Data Fig. 2.**
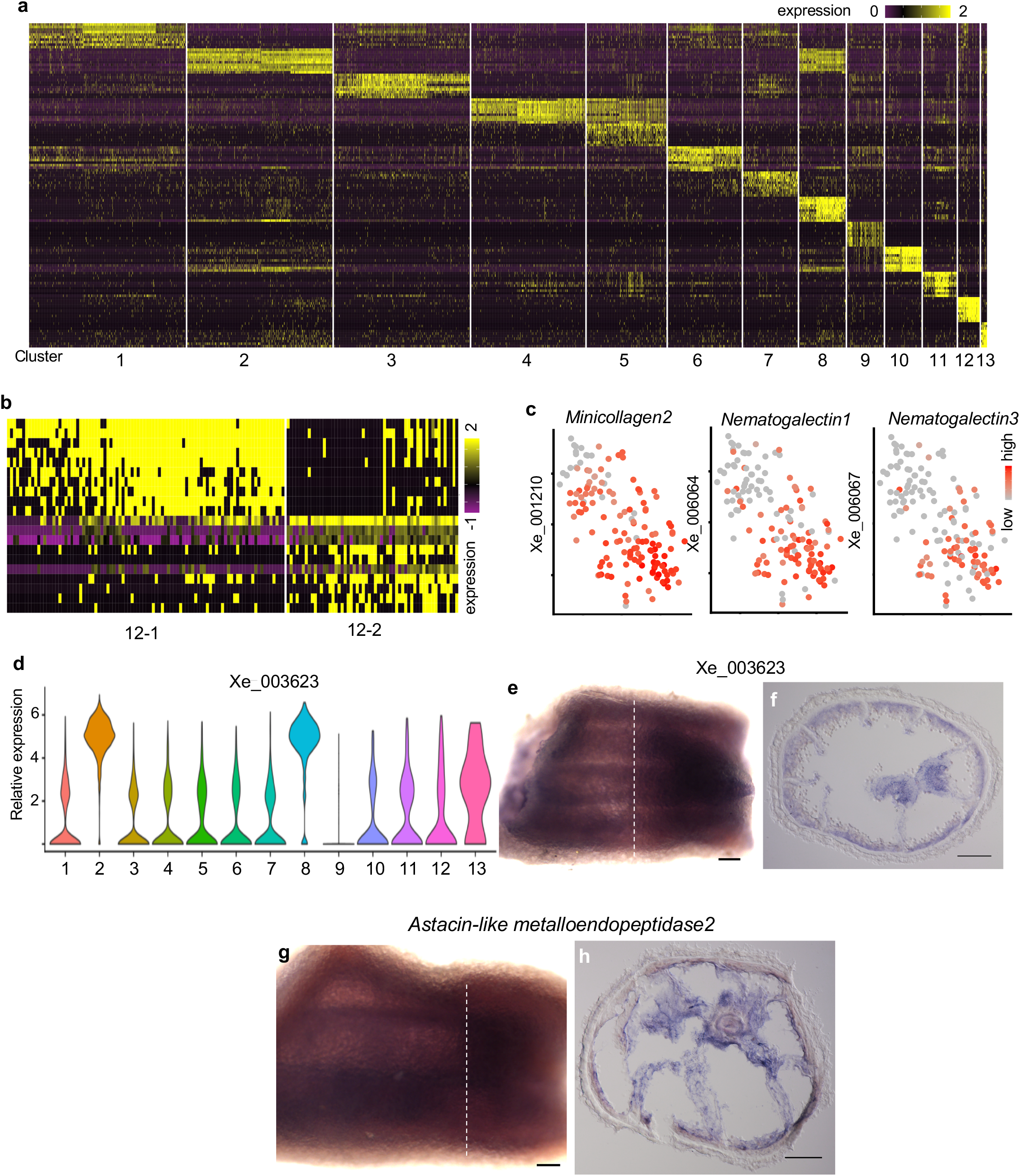
Additional scRNA-seq analyses. **a**, Heatmap showing differential gene expression patterns of all cells in the assigned 13 cell clusters (indicated at the bottom). Each column is one cell cluster and each row represents one gene. **b**, Heatmap showing differential gene expression patterns of two sub-clusters in cluster 12, 12-1 and 12-2. **c**, Expression levels (see colored expression scale) of 3 cluster 12 markers, *Minicollagen2*, *Nematogalectin1*, and *Nematogalectin3* are shown in the tSNE plots. **d**, Expression levels of Xe_003623, a non-conserved and uncharacterized cluster 13 marker genes, in each of the 13 cell types defined by scRNA-seq and Seurat. **e-h**, RNA *in situ* hybridization of Xe_003623 (**e, f**) and *Astacin-like metalloendopeptidase2* (**g, h**). The white dashed lines in the whole mount views (**e,g**) indicate the cross section cut (**f, h**). Scale bars, 150 μm (**e**, **g**), 100 μm (**f**, **h**)

**Extended Data Fig. 3.**
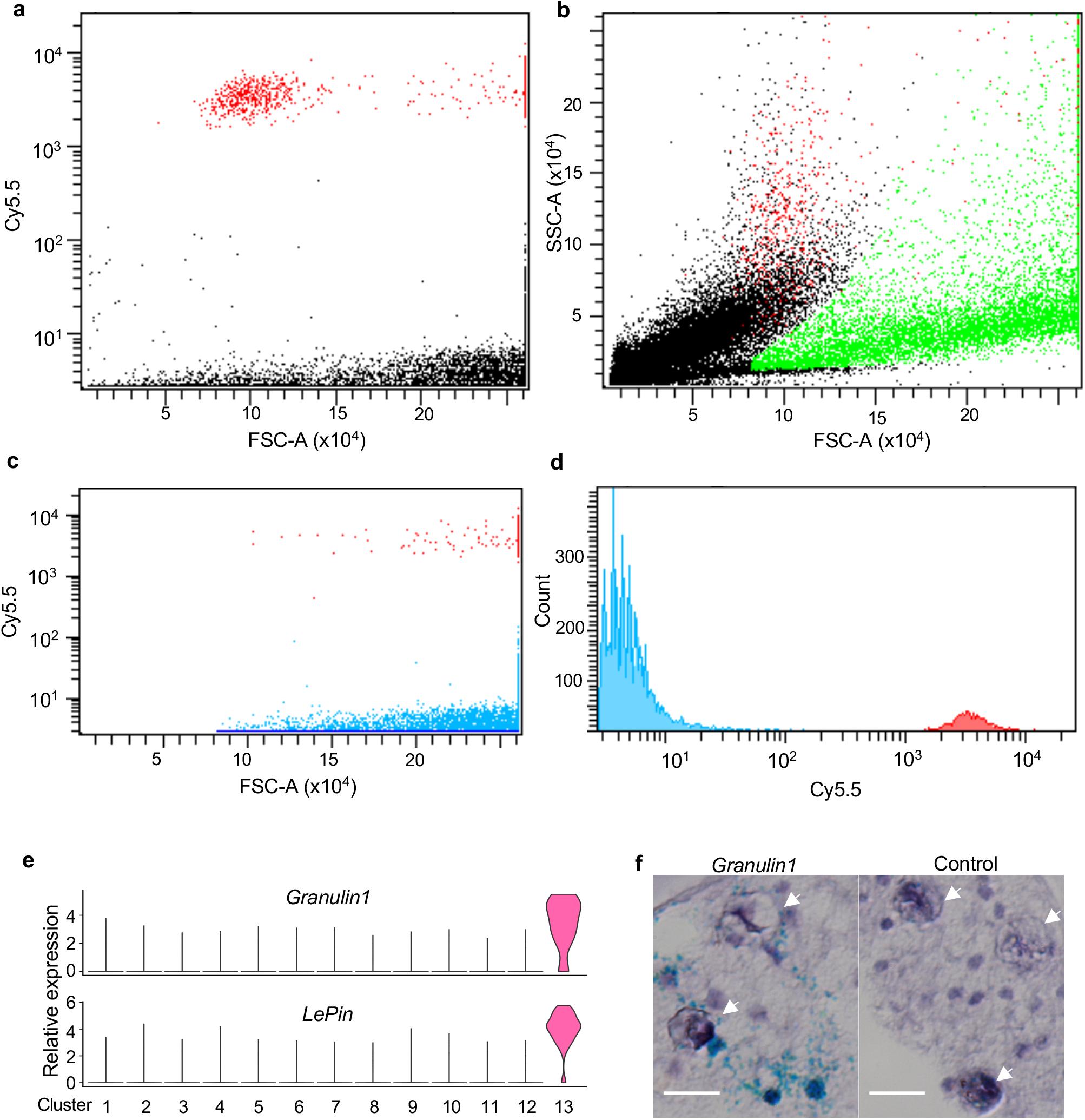
Additional FACS and scRNA-seq analyses. **a**, Dissociated *Xenia* cells were first sorted by Cy5.5 and FSC-A to identify the potential algae-containing cells (Red dots). **b**, Mapping the potential algae-containing population (red dots) onto the FSC-A and SSC-A space suggests that these are real cells. To gate for algae-free cells, we first used DAPI to stain the nuclei of dissociated, fixed, and permeabilized *Xenia* cells and identified the DAPI positive sub-population (green dots), which represents cells. **c**. Further removing of Cy5.5 strong algae-containing cells among the DAPI positive population led to the sorting of algae-free coral cells (blue dots). **d**, The counts distribution of algae-containing (red) and algae-free (blue) cells in our sorting. **e**, Piano plots of the expression profiles of *Granulin1* and *LePin* in the 13 clusters defined by scRNA-seq and Seurat. **f**, Ultra-sensitive chromogenic RNA *in situ* hybridization by RNAscope probing for *Granulin1* (left) and control (right). Positive signals are blue. Samples were countered stained with Hematoxylin. White arrows indicate *Symbiodiniaceae*. Scale bars, 10 μm.

**Extended Data Fig. 4.**
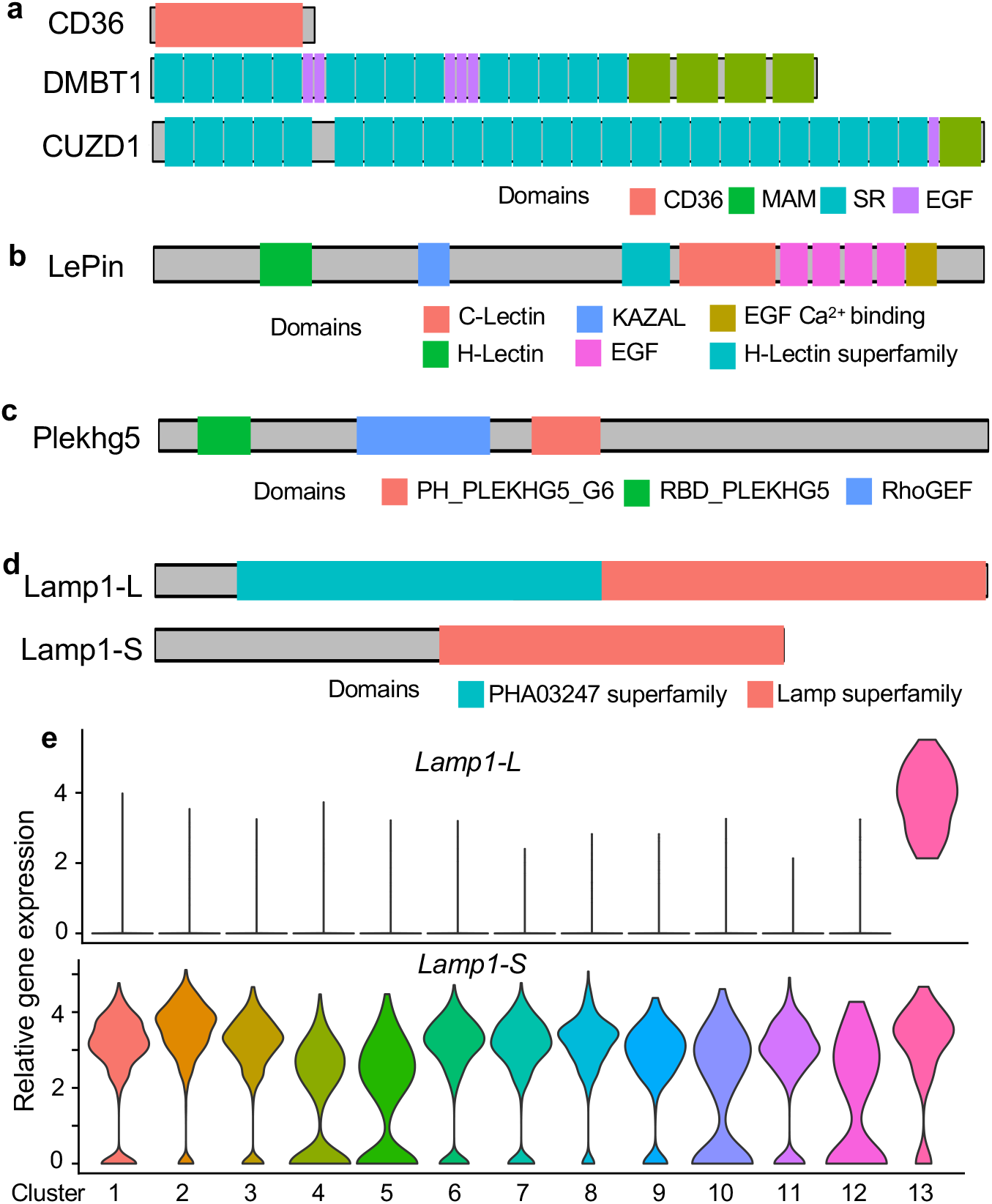
Selected endosymbiotic marker genes with known domains. **a**, Scavenger receptors: CD36, DMBT1, and CUZD1. **b**, LePin. **c**, Plekhg5. **d**, Lamp1-L and Lamp1-S. **e**, Piano plots of expression profiles of Lamp1-L and Lamp1-S in 13 clusters defined by scRNA-seq and Seurat.

**Extended Data Fig. 5.**
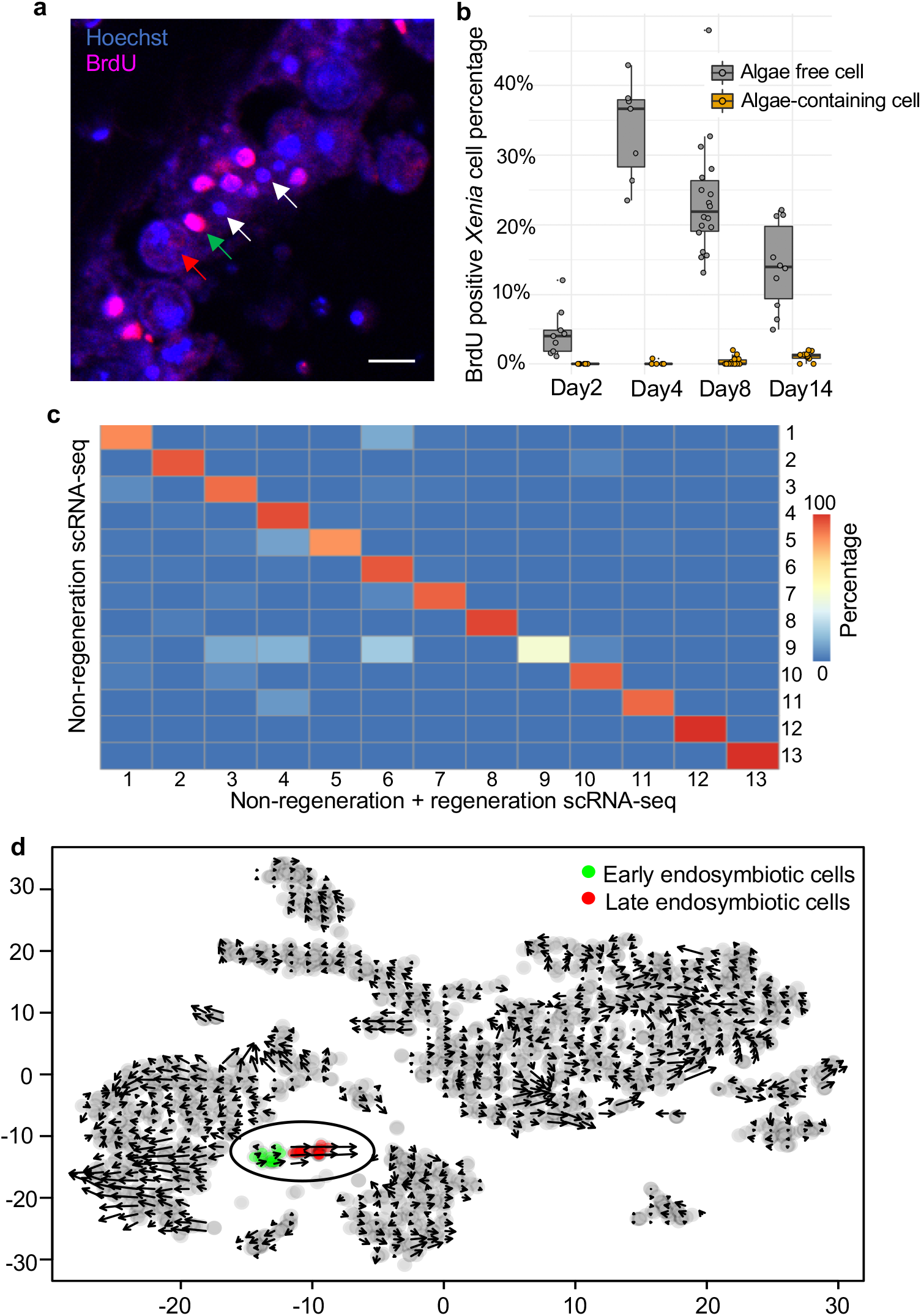
Additional analyses of endosymbiotic cell lineage. **a,** A representative image of BrdU labeling (red), overlaid with Hoechst (blue DNA stain) in a cross section of a regenerating *Xenia sp*. stalk. White, red, and green arrows indicate BrdU-negative (BrdU^−^) *Xenia* nuclei, an alga, and a BrdU-positive (BrdU^+^) *Xenia* nucleus juxtaposed to the alga, respectively. **b**, Boxplot. Percentages (y-axis) of BrdU^+^ *Xenia* cells at the indicated regeneration time points (x-axis). Each dot represents data from one section. Three independent samples were pooled and plotted for each time point. The algae-containing proliferated *Xenia* cells were estimated as those whose BrdU^+^ nuclei were juxtaposed to algae. The medians are indicated as lines in the box. The upper and lower edge of the box represents upper and lower quartiles, respectively. **c**, Comparisons of cell clusters between the non-regeneration samples (y-axis) and all samples (i.e. regeneration and non-regeneration samples combined, x-axis). The heatmap shows the percentage of cells in each of the 13 cluster defined by the non-regeneration samples that are found in the corresponding 13 types defined by all samples. **d**, Velocyto analysis of the scRNA-seq data from day 4 regenerating *Xenia sp*. stalks. Each dot represents a cell and arrows indicate the directions of RNA-velocity. The endosymbiotic cell cluster is outlined and the predicted early and late cell states are colored green and red, respectively.

**Extended Data Table 1.** Summary of Illumina sequencing for genome assembly. The table shows sequence information from 4 library preparations for Illumina sequencing. Read number indicates total number of reads obtained. Read length indicates individual read length, by paired-end sequencing. Data indicate total sequence data in giga bases.

| library | read number (M*) | read length | pair-end | data (G**) |
| --- | --- | --- | --- | --- |
| 1 | 27.2 | 150 | No | 4.09 |
| 2 | 19.7 | 150 | No | 2.91 |
| 3 | 73.9 | 75 | No | 5.54 |
| 4 | 127.4 | 150 | Yes | 38.2 |

\* M, million; \*\* G, giga base

**Extended Data Table 2.** Summary of Nanopore sequencing for genome assembly. Statistics of all sequence information from 4 different runs of Nanopore sequencing, including max, mean, and median reads length statistics, and the percentages of reads that have bigger sizes than the indicated number, > 5 kb, > 10 kb, and > 20 kb.

| library | read number | max (bp*) | mean (bp) | median (bp) | > 5 kb** | >10 kb | >20 kb | data (G***) |
| --- | --- | --- | --- | --- | --- | --- | --- | --- |
| run1 | 819,205 | 62,624 | 4,488 | 4,341 | 38.4% | 3.5% | 0.19% | 3.68 |
| run2 | 328,045 | 266,931 | 11,716 | 4,343 | 46.1% | 29.9% | 17.3% | 3.84 |
| run3 | 210,985 | 310,074 | 12,677 | 5,400 | 52.4% | 32.7% | 18.3% | 2.67 |
| run4 | 1,912,621 | 143,219 | 5,228 | 4,959 | 49.5% | 7.27% | 0.37% | 10 |

\* bp, base pair; \*\* kb, kilobase; \*\*\* G, giga base

**Extended Data Table 3.** Transcriptomes for gene annotation. A summary of all the transcriptome data used for gene annotation. RNA isolated from different samples as indicated were used for Illumina sequencing in order to cover as many expressed gene as possible. Opti-Prep, density-based separation of dissociated *Xenia* cells into 4 different layers (see Method). lv1, lv2, lv3, and lv4 indicate layer 1, layer 2, layer 3, and layer 4 cells, respectively, used to make the RNA-seq libraries.

| samples | library | read number (M*) | data (G**) |
| --- | --- | --- | --- |
| whole polyp | pair-end | 34.2 | 5.12 |
| Stalk | pair-end | 36.0 | 5.40 |
| Tentacle | pair-end | 39.8 | 5.97 |
| Regeneration 4d stalk | pair-end | 38.9 | 5.84 |
| Opti-Prep lv1 | pair-end | 29.6 | 4.43 |
| Opti-Prep lv2 | pair-end | 31.8 | 4.76 |
| Opti-Prep lv3 | pair-end | 33.6 | 5.03 |
| Opti-Prep lv4 | pair-end | 31.1 | 4.68 |

\* M, million; \*\* G, giga base

**Extended Data Table 4.** Comparisons of *Xenia sp*. genome assembly with the indicated published cnidarian genomes. Number of gene models indicate predicted gene model number, whereas number of complete gene models represent number of genes with clearly predicted in-frame start and stop codons.

| Parameter | species | *Xenia*<br>*sp.* | *Aiptasia* | *Nematostella*<br>*vectensis* | *Acropora*<br>*digitifera* |
| --- | --- | --- | --- | --- | --- |
| Predicted genome size (Mb) |  | 197 | 260 | 329 | 420 |
| Assembly size (Mb) |  | 222.7 | 258 | 356 | 419 |
| Total contig size (Mb) |  | 222.5 | 213 | 297 | 365 |
| Total contig size as % of assembly size |  | 99.9 | 82.5 | 83.4 | 87 |
| Contig N50 (Kb) |  | 1,122 | 14.9 | 19.8 | 10.9 |
| Scaffold N50 (Kb) |  | 148,322 | 440 | 472 | 191 |
| Number of gene models |  | 29015 | 29269 | 27273 | 23668 |
| Number of complete gene models |  | 27640 | 26658 | 13343 | 16434 |
| Mean exon length (bp) |  | 204 | 354 | 208 | 230 |
| Mean intron length (bp) |  | 448 | 638 | 800 | 952 |
| Mean protein length (number of amino acids) |  | 414 | 517 | 331 | 424 |

## Coral aquarium

The coral aquarium is established in a tank (Reefer 450 system, Red Sea). The artificial sea water made from Coral Pro Salt (Red Sea) was first incubated with live rocks for two months before introducing *Xenia sp*., other corals, fish, snails, and hermit crabs. The aquarium is maintained at ~80°F with ~25% change of sea water every 1-2 weeks. The light is provided by Hydra 26™ HD LED (Aqua Illumination) with 60% power on during 10:00 AM to 7:30PM. The fish were fed with fish pellets (New Life Spectrum Marine Fish Formula) and Green Marine Algae (Ocean Nutrition).

## Genomic DNA isolation from *Xenia sp.*

To enable Nanopore DNA sequencing, we modified a protocol^31^ that allowed the isolation of long DNA fragments. For each DNA preparation, one or two *Xenia sp*. colonies containing ~30 polyps were collected from the aquatic tank and washed 3 times for 5 min each with Ca^2+^ and Mg^2+^ free artificial sea water (449 mM NaCl, 9 mM KCl, 33 mM Na_2_SO_4_, 2.15 mM NaHCO_3_, 10 mM Tris-HCl, 2.5 mM EGTA, pH8.0). Tentacles were cut away as the mucus surrounding them affected the quality of the isolated DNA. The remaining stalks and the bases of individual *Xenia* colonies were placed in 100 μl DNAzol (Invitrogen) in a 1.5 ml microfuge tube. The tissues were cut into small pieces by a scissor to make the fragment sizes of ~1/10^th^ of the original size. These fragments were further minced by a small pestle made for 1.5 ml microfuge tubes (Fisher Scientific, 12-141-364). Then 900 μl DNAzol were added followed by vortexing the sample and then transferred to a 15 ml conical tube. 4 ml of DNAzol and 50 μl of 10 mg/ml RNase A were then added to the tube and mixed followed by incubation at 37°C for 10 min. 25 μl of 20 mg/ml proteinase K were then added, mixed, and the tube was incubated at 37°C for another 10 min. The sample was centrifuged at 5000xg for 10 min. The supernatant was transferred to another 15 ml tube. After adding 2.5 ml ethanol, the tube was gently mixed by inverting several times. The tube was let to stand at room temperature for 3 min followed by centrifugation at 1000xg for 10 min to pellet the DNA. The supernatant was discarded and the DNA pellet was resuspended in with 500 μl TE (10 mM Tris-HCl, 1 mM disodium EDTA, pH 8.0) buffer. After the DNA had dissolved, 500 μl of phenol: chloroform: isoamyl alcohol (25:24:1) was added, and the tubed was placed on the Intelli-Mixer RM-2S for mixing using program C1 at 35 rpm for 10 min. The mixture was then transferred to a 2 ml phase-lock gel (QuantaBio, Cat. 2302820) and centrifuged at 4500 rpm for 10 min. The aqueous phase was transferred into a new 2 ml tube, 200 μl 5 M ammonium acetate and 1.5 ml ice-cold ethanol were added followed by centrifugation at 10,000xg for 10 min to pellet DNA. The pellet was washed twice with 1 ml 80% ethanol. After removing as much ethanol as possible, the DNA pellet was left to dry at 42°C for 1 min, and then resuspended in 50 μl TE buffer.

## Illumina sequencing

Genomic DNA prepared as above was fragmented into ~400 bp and libraries were made with ThruPLEX DNA-Seq kit (TaKaRa) according to the manufacturer’s manual. These libraries were sequenced using the NEXseq500 platform with NextSeq^®^ 500/550 High Output Reagent Cartridge v2 (Illumina).

## Nanopore sequencing

Genomic DNA was used to build Nanopore sequencing libraries with Ligation Sequencing Kit (SQK-LSK108, Oxford Nanopore Technologies), following the manufacturer’s manual. For the first 3 runs, genomic DNA was not fragmented in order to generate long reads. To obtain more reads, for the 4^th^ run of Nanopore sequencing, genomic DNA was sheared to 8-10 kilobases by g-TUBE™ (Covaris, 520079). The libraries were sequenced in R9.4.1 flow cells on MinION device (Oxford Nanopore Technologies) and the data were combined for genome assembly.

## Chromosome conformation capture (Hi-C)

To perform Hi-C on *Xenia sp*. tissue, we modified our previously published protocol for nuclear in situ ligation^32^ as described in detail below.

### 1. Fix and dissociate tissues

Fix 8 polyps (~10^8^ cell) with 4% paraformaldehyde (PFA) overnight. After washing twice with 3.3x PBS^33^ and dissociating the tissue in 2 ml 3.3x PBS using a 7 ml glass Dounce tissue grinder (Wheaton), another 3 ml 3.3x PBS was added. The mixture was then transfer to a 15 ml conical tube and centrifuged at 1000g for 3 min (Sorvall Lynx 6000 centrifuge, ThermoFisher Scientific). The pellet was washed once with 5 ml 3.3x PBS.

### 2. Nuclear permeabilization and chromatin digestion

The pellet from step 1 was resuspended in 10 ml ice-cold Hi-C lysis buffer (10 mM Tris, pH8.0, 10 mM NaCl, 0.2% NP-40, 1x protease inhibitors cocktail (Roche, 04693132001)) and rotated for 30 min at 4°C followed by centrifugation at 1,000g for 5 min at 4°C. The pellet was washed with 1 ml ice-cold 1.2x NEB3.1 (120μl NEB3.1 to 880μl ddH_2_O) buffer and transferred to 1.5 ml microfuge tube followed by centrifugation at 1,000g for 5 min at 4°C. The pellet was washed again with 1 ml ice-cold 1.2x NEB3.1 followed by centrifugation. After removing the supernatant, 400 μl 1.2x NEB3.1 buffer and 12 μl of 10% SDS were added to the pellet. P200 pipet tip was used to thoroughly resuspend and dissociate the pellet. The mixture was then incubated at 65°C for 10 min at 950 rpm in a Thermomixer (Eppendorf). After cooling the mix on ice for 5 min, 40 μl 20% Triton X-100 was added to the mixture to neutralize the SDS. After carefully mixing by pipetting with a P200 pipet tip and inverting the tube several times, the mixture was then incubated at 37°C for 60 min with rotation (950 rpm) in a Thermomixer. To digest the crosslinked genomic DNA, 30 μl of 50 U/μl BglII (NEB R0144M) was added to the mixture and incubated overnight at 37°C with rotation at 950 rpm in a Thermomixer.

### 3. Fill in 5’overhang generated by BglII digestion with biotin

A nucleotide mix containing dATP, dGTP and dTTP was made by adding 1μl each of 100 mM dATP, dGTP and dTTP into 27μl ddH2O. To the 480.0μl BglII digested nuclear preparation from the above step 2, 4.5 μl of the nucleotide mix, 15 μl 1 mM biotin-16-dCTP (Axxora, JBS-NU-809-BIO16) and 10 μl 5U/μl Klenow (NEB, M0210L) were added followed by incubation at 37°C for 90 min with intermittent gentle shaking at 700 rpm for 10s after every 20s using Thermomixer. The tube was also taken out and inverted every 15-20 min. After this incubation, the mixture was kept on ice.

### 4. Proximity ligation

The mixture from step 3 was transferred to a 50 ml conical tube followed by adding 750 μl 10x T4 ligase buffer (NEB B0202S, no PEG), 75 μl 100x BSA (NEB), 6140.5 μl water, 25 μl 30 U/μl T4 DNA ligase (Thermo Scientific, EL0013), and incubating at 16°C overnight.

### 5. Reverse crosslink and DNA isolation

25 μl 20 mg/ml proteinase K (Invitrogen, 25530-049) was added into the reaction mixture from step 4 and the mixture was divided equally into 8×1.5 ml microfuge tubes (~950 μl per tube). The tubes were then incubated overnight at 65°C with rotation at 950 rpm in a Thermomixer. Next day, 3 μl 20 mg/ml proteinase K were added to each tube followed by incubation at 65°C for 2 hours with mixing in Thermomixer. The mixtures were combined into one 50 ml conical tube. After cooling down to room temperature, 10 ml phenol (pH 8.0) (Sigma) was added and mixed by vortex for 2 min. The mixture was then centrifuged for 10 min at 3,000g (Sorvall Lynx 6000 centrifuge). The supernatant containing the DNA was mixed with 10 ml phenol:chloroform (1:1) (pre-warmed to room temperature) and vortexed for 2 min. The whole mixture was then transferred to a 50 ml MaXtract High Density tube (Qiagen, 129073) and centrifuged at 1,500g for 5 min (Sorvall Lynx 6000 centrifuge). The top phase containing the Hi-C DNA was transferred to a 50 ml conical tube and the volume (usually ~10 ml) was adjusted to 10 ml with 1xTE as needed. To pellet the DNA, 1 ml 3M Na-acetate, 5 μl 15 mg/ml GlycoBlue (Invitrogen AM9515) and 10 ml isopropanol were added to the mixture and incubated at −80°C for >1 hour. The DNA was then pelleted by centrifugation at 17,000g for 45 min at 4°C (Sorvall Lynx 6000 centrifuge). The Hi-C DNA pellet was resuspended in 450 μl 1xTE and transferred to an 1.5 ml microfuge tube followed by adding 500 μl phenol:chloroform (1:1). After mixing by vortex, the mix was centrifuged at 18,000g for 5 min at room temperature. The top aqueous layer was collected into another tube followed by adding 40 μl 3M Na-acetate, 1 μl 15 mg/ml GlycoBlue (Invitrogen AM9515, 300μl) and 1 ml ice-cold 100% ethanol. After incubating at −80°C for >30 min, the DNA was centrifuged at 21000g for 30 min at 4°C. The DNA pellet was washed with freshly prepared 70% ethanol and air dry followed by dissolving in 45 μl EB (10mM Tris, pH8.0). The contaminated RNA in the DNA preparation was digested by adding 0.5 μl 10 mg/ml RNaseA and incubated at 37°C for 30 min.

### 6. Remove biotin from the free DNA (un-ligated DNA) ends

To remove the biotin at the free DNA ends, 1.0 μl 10 mg/ml BSA (NEB, 100x), 10.0 μl 10x NEB 2.1 buffer, 1 μl 10 mM dATP, 1 μl 10 mM dGTP and 5 μl T4 DNA polymerase (NEB M0203S), and 42 μl water were added to 40 μl (~ 3μg) Hi-C DNA preparation from step 7. The mixture was divided into two equal aliquots in 2 PCR tubes and incubate at 20°C for 4 hours. 2 μl 0.5 M EDTA was added to each of the two tubes to stop the reason. The Hi-C DNA was then purified using the Clean and Concentrator Kit (ZYMO, D4013) followed by elution with 50 μl EB.

### 7. Biotin pull down of DNA and second DNA digestion

60 μl Dynabeads MyOne Streptavidin C1 (Invitrogen) was washed in 1.5 ml non-sticking microfuge tubes (Ambion) with 200 μl 2x binding buffer (BB, 10 mM Tris, pH8, 0,1 mM EDTA, 2 M NaCl) twice followed by resuspension in 50 μl 2x BB. The 50 μl Hi-C DNA from step 6 was added followed by rotating for 30 min using Intelli-Mixer (ELMI) at room temperature. The beads were collected using a magnetic stand and washed with 100 μl 1x BB followed by washing with 100 μl 1x NEB4 buffer twice and resuspending in 50 μl 1x NEB4 buffer. The DNA on beads was digested using 1 μl 10U/μl AluI (NEB, R0137S) at 37°C for 60 min. The beads were collected on a magnetic stand followed by washing with 100 μl 1x BB and then 100 μl EB. The beads were resuspended in 30 μl EB.

### 8. A-tailing

The 30 μl beads from step 7 were mixed with 5 μl NEB Buffer 2, 10 μl 1 mM dATP, 2 μl H_2_O, 3 μl Klenow (3’-5’ exo-) (NEB M0212L) and incubated at 37°C for 45 min. After the reaction, the beads were collected by a magnetic stand followed by washing with 100 μl 1x BB and then 100 μl EB. The beads were resuspended in 50 μl EB.

### 9. Sequencing adaptor ligation

The 50 μl beads from step 8 was mixed with 3.75 μl sequencing adapter (TruSeq RNA Sample Prep Kit v2), 10 μl 1x T4 DNA ligase buffer, 3 μl T4 DNA Ligase (30U/μl) (Thermo Scientific, EL0013) and incubate at room temperature for 2 hours. The beads were collected by a magnetic stand followed by washing twice with 400 μl 1x BB + 0.05% Tween, 200 μl 1x BB, and then 100 μl EB. The beads were resuspended in 40 μl EB. To release the DNA from the beads, the mixture was incubated at 98°C for 10 min and then centrifuged at 500 rpm to pellet the streptavidin beads.

### 10. TruSeq RNA Library Prep Kit was used to make DNA sequencing library (8 PCR cycle was used) and the DNA was sequenced by NextSeq 500

## Single Cell RNA-seq (scRNA-seq)

1 polyp, 8 tentacles, two stalks, or 2 regenerating stalks of *Xenia sp*. were dissociated into single cells in 1 ml digestion buffer, containing 3.6 mg/ml dispase II (Sigma, D4693), 0.25 mg/ml Liberase (Sigma, 5401119001), 4% L-cysteine in Ca^2+^ free sea water (393.1 mM NaCl, 10.2 mM KCl, 15.7 mM MgSO_4_·7H_2_O, 51.4 mM MgCl_2_·6H_2_O, 21.1 mM Na_2_SO_4_, and 3 mM NaHCO_3,_ pH 8.5) and incubated for 1 hour at room temperature. After digestion, fetal bovine serum was added to a final concentration of 8% to stop enzymatic digestion. Cell suspension was filtered through a 40 μM cell strainer (FALCON). Cells were counted by hemocytometer and 17000 cells per tissue sample were subjected to single cell library preparation using the 10x Genomics platform with Chromium Single Cell 3’ Library and Gel Bead Kit v2 (PN-120267), Single Cell 3′ A Chip Kit (PN-1000009), and i7 Multiplex Kit (PN-120262). Libraries were sequenced using Illumina NextSeq 500 for paired-end reads. Read1 is 26 bp while Read2 is 98 bp.

## Bulk RNA-seq

Total RNA was isolated from 3 polyps, 32 tentacles, or 6 stalks by RNeasy Plus Mini Kit (QIAGEN). To obtain additional transcriptome from different cell types, we dissociated coral tissue into individual cells according to a previously published method^34^ and subjected the dissociated cells to OptiPrep based cell separation^35^. Cells with different density were separated into four different layers and RNA was isolated from each layer with the same kit described above. For transcriptome of FACS sorted algae-containing and algae-free cells, 3 polyps were dissociated with the same protocol as used in the scRNA-seq and the dissociated cells were subjected to FACS. Cy5.5 positive and negative cells were collected as algae-containing and algae-free cells, respectively, and used for total RNA extraction as above. cDNA libraries were built according to TruSeq Stranded mRNA Library Prep Kit (Illumina) and subjected to Illumina NextSeq 500 for sequencing. For gene annotation, paired-end sequencing of 75 bp for each end was used. For FACS sorted bulk cell transcriptomes, single end sequencing of 75 bp was used.

## *Xenia* regeneration and BrdU tracing

Individual *Xenia sp*. polyps were placed into a well of 12-well cell culture plate (Corning) containing 2 ml artificial sea water from the aquatic tank. The polyps were allowed to settle in the well for 5~7 days before cutting away the tentacles. For BrdU tracing, 0.5 mg/ml BrdU was added into the well 2 days prior to sample harvest. The BrdU labeled stalks were fixed by 4% PFA overnight followed by washing with PBST (PBS+0.1% Tween 20) twice for 10 min each. The stalk was then balanced with 30% sucrose overnight followed by embedding in OCT, frozen in dry ice/ethonal and subjected to cryo-sectioning. The slides were washed with PBS 3 times for 5 min each time followed by treating with 2 M HCl containing 0.5% Triton X-100 for 30 min at room temperature. The slides were then incubated with PBST (0.2% Triton X-100 in PBS) 5 min for 3 times each followed by blocking with 10% goat serum and then incubating with mouse anti-BrdU antibody (ZYMED, 18-0103, 1:200 dillution in 10% goat serum) at 4°C overnight. Slides were washed with PBST 3 times for 10 min each followed by incubation with the secondary antibody (Invitrogen) for 1 hour at room temperature and washing with PBST 3 times for 10 min each. The nuclei were counterstained with Hoechst 33342 and the signal was visualized using a confocal microscope (Leica). Clear BrdU signal in the nucleus labeled by Hoechst was counted as a BrdU^+^ cell. If the *Xenia* BrdU^+^ nucleus was juxtaposed to an alga, it was counted as an alga containing BrdU^+^ *Xenia* cell.

## Whole mount RNA *in situ* hybridization

To perform RNA *in situ* hybridization on *Xenia*, we modified the whole mount RNA *in situ* hybridization protocol for zebrafish^36^ as described below.

For making gene specific probes, we design primers (Supplementary table 6) to genes of interest for polymerase chain reaction (PCR) to amplify gene fragments from *Xenia sp*. cDNA. T3 promoter sequence was added to the 5’ of the reverse primers so that the PCR products could be directly used for synthesizing anti-sense RNA probes by T3 RNA polymerase (Promega, P2083) using DIG RNA Labeling Mix (Roche, 11277073910). DIG-labeled RNA probes were purified by RNA Clean and Concentrator-5 (ZYMO), heated to 80°C for 10 min, immediately transferred on ice for 1 min, and then diluted in Prehyb^+^ buffer (50% Formamide, 5XSSC, 50 μg/ml Heparin, 2.5% Tween 20, 50 μg/ml SSDNA (Sigma, D1626)) to a final concentration of 0.5 μg/ml, and stored in −20°C until use.

*Xenia* polyps were relaxed in Ca^2+^ free sea water (described above) for 30 min and fixed in 4% PFA in Ca^2+^ free sea water overnight at 4°C. Fixed polyps were washed with PBST (0.1% Tween 20 in PBS) twice for 10 min each, and then incubated in 100% methanol at −20°C overnight. Next day, the tissues were washed sequentially in 75%, 50%, and 25% methanol for 5 min each and then washed in PBST for 10 min. They were then treated with 50 μg/ml proteinase K in PBST for 20 min followed by washing in PBST briefly. The tissues were post-fixed in 4% PFA at room temperature for 20 min and then washed with PBST 2 time for 10 min each. Pre-hybridization was performed in Prehyb^+^ at 68°C for 2 h, followed by incubation with probes in Prehyb^+^ overnight at 68°C. After probes were removed, samples were washed sequentially in 2X SSC (0.3 M NaCl and 0.03 M sodium citrate) containing 50% formamide for 20 min twice, 2X SSC containing 25% formamide for 20 min, 2X SSC for 20 min twice, and 0.2X SSC for 30 min 3 times each, all at 68°C. Then, samples were washed in PBST at room temperature for 10 min and incubated in DIG blocking buffer (1% ISH blocking reagent (Roche, 11096176001) in maleic acid buffer (0.1M maleic acid, 0.15 M NaCl, pH 7.5) for 1 hour at room temperature, followed by incubation in anti-DIG antibody (Anti-Digoxigenin-AP (Roche, 11093274910)) at 1:5000 dilution in DIG blocking buffer overnight at 4°C. Next day, the samples were wash in PBST for 10 min 3 times each at room temperature, then in 9.5T buffer (100 mM Tris-HCl pH9.5, 50 mM MgCl_2_, 100 mM NaCl, 0.1% Tween 20) for 10 min 3 times each at room temperature. Hybridization signals were revealed by incubation in BCIP/NBT buffer (1 SIGMAFAST™ BCIP^®^/NBT tablet (Sigma, B5655) in 10 ml H_2_O)) at 4°C until brown-purplish colors were sufficiently dark. For this study, the color development step took 48-60 hours. The samples were then wash in PBST twice for 10 min each. The samples were post-fixed in 4% PFA overnight at 4°C followed by washing in PBST twice for 10 min each and then washing in methanol for 3 h at room temperature. The tissues were kept in PBS and imaged using SMZ1500 microscope (Nikon) under Ring Light System (Fiber-Lite). For cross section of stalks, the whole mount sample was processed for frozen cryosection as described above.

## RNAscope *in situ* hybridization assay for *LePin* and *Granulin1* expression

To visualize RNA expression in endosymbiotic cells, we used the ultrasensitive RNAscope ISH approach (Advanced Cell Diagnostics Inc, ACD). *LePin*- *or Granulin1-*specific oligo probes were ordered from ACD (see Supplementary Table S6 for further information). The fluorescent RNAscope assay was carried out by RNAscope® Multiplex Fluorescent Reagent Kit v2 (ACD) according to manufacturer’s protocol. The Chromogenic assay was carried by RNAscope® 2.5 HD Duplex Detection Kit (ACD) according to manufacturer’s protocol. Both assays used the cryo-section of the fixed *Xenia* polyp prepared according to the manufacturer’s protocol.

## Genome assembly

Sequencing data from Nanopore were used to initiate the genome assemble by Canu (v1.7)^37^. The assembled genome was further polished with Illumina short reads by Nanopolish (v0.9.2, https://github.com/jts/nanopolish) with 5 cycles, which resulted in 1482 high quality contigs for the diploid genome. The diploid genome assembly was separated into haploid by HaploMerger2^38^. The haploid genome assembly was further subject to Hi-C assisted scaffolds by 3D *de novo* assembly pipeline^39^.

## Gene annotation

Funannotate genome annotation pipeline (v1.3.3, https://github.com/nextgenusfs/funannotate) was used to annotate the *Xenia sp*. genome. In brief, transcriptome data were assembled by Trinity (v2.6.6)^40^ and used to generate the gene models based on the presence of mRNA by PASApipeline (v2.3.2)^41^. These gene models were used as training sets to perform de novo gene prediction by AUGUSTUS (v3.2.3)^42^ and GeneMark-ES Suite (v4.32)^43^. All gene models predicted by PASApipeline, AUGUSTUS, and GeneMark were combined and subject to EVidenceModeler to generate combined gene models^44^. The predicted genes were filtered out if more than 90% of the sequence overlapped with repeat elements as identified by RepeatMasker and RepeatModeler (http://www.repeatmasker.org). PASA was further used to add 3’ and 5’ untranslated region (UTR) sequences to the remaining predicted genes. Pfam (v31.0), Interpro (v 67.0), Uniprot (v2018_03), BUSCO (v1.0)^45^ databases and eggnog-mapper (v1.3)^46^ were used to annotate the function of these gene models.

## Phylogeny tree analysis

OrthoFinder (v2.2.7)^47^ was used to find orthologs from different species and build the phylogenetic tree. Diamond (v0.9.21) is used to align the orthologs. MSA and FastTree methods were used to infer the gene trees and phylogenetic tree.

## Single cell clustering and marker gene identification

The raw single cell sequencing data was de-multiplexed and converted to FASTAQ format by Illumina bcl2fastaq software. Cell Ranger (v2.1.1, https://support.10xgenomics.com/single-cell-vdj/software/pipelines/latest/what-is-cell-ranger) was used to de-multiplex samples, process barcodes and count gene expression. The sequence was aligned to the annotated *Xenia sp*. genome and only the confidently mapped and non-PCR duplicated reads were used to generate gene expression matrix. Cells with UMI numbers less than 400 or mitochondria gene expression > 0.2% of total genes were filtered out. To further remove outliers, we calculated the UMI number distribution detected per cell and removed cells in the top 1% quantile. To remove batch effect, we applied Seurat canonical correlation analysis (CCA) alignment method for data integration^48^. For each dataset, we identified top 1000 genes with the highest dispersion. We combined the top 1000 genes from each dataset and ran a canonical correlation analysis to generate the CCA space. The first 16 canonical correlation vectors were used to generate the dimensional reduction which was used for further clustering analysis. Clustering and marker gene identification was performed with Seurat by setting the resolution to 0.6, which led to the identification of 13 cell clusters.

## Identification of *Xenia sp*. cells performing endosymbiosis with *Symbiodiniaceae*

The bulk transcriptome data of FACS isolated algae-containing or algae-free cells were aligned to *Xenia sp*. genome by STAR (v2.5.3a)^49^. Individual gene expression (RPKM) for each sample were calculated by RSEM (v1.3.0)^50^. The gene expression levels of each bulk RNA-seq of FACS-sorted cells were compared with the gene expression levels calculated using average UMI (unique molecular index) number for each gene in each cell cluster identified by scRNA-seq. Pearson correlation coefficient was calculated for each comparison.

## Pseudotime analysis

To infer the trajectory of endosymbiotic *Xenia* cells, we combined scRNA-seq data of regenerating and non-regenerating samples by Seurat CCA alignment. All cells belonging to the endosymbiotic cell cluster (Cluster 13) were subjected to Monocle (v2.10.1)^51^ analyses. These cells were further divided into two sub-clusters and the top 1000 differentially expressed genes between these two groups were used as ordering genes to construct the trajectory by DDRTree algorithm. The differentially expressed genes along pseudotime is detected by differentialGeneTest function in Monocle.

## RNA velocity

RNA velocity estimation was carried out by velocyto.R program (http://velocyto.org, v0.6) according to the instruction^52^. Briefly, velocyto used raw data of regeneration sample to count the spliced (mRNA) and un-spliced intron reads for each gene to generate a loom file. This loom file was loaded into R (v3.5) by read.loom.matrices function and used to generate RNA velocity map. The RNA velocity map was projected into the same t-SNE space identified by Seurat.

## Data availability

Raw sequence data for this study is available in NCBI BioProject under accession PRJNA548325. Assembled genome and annotation files are available at https://drive.google.com/open?id=1qmqYMepjIsYnOrBuUAe_1-GB-y8oqy0C.

